# Deletion mapping of regulatory elements for *GATA3* reveals a distal T helper 2 cell enhancer involved in allergic diseases

**DOI:** 10.1101/2022.05.24.493112

**Authors:** Hsiuyi V. Chen, Patrick C. Fiaux, Arko Sen, Ishika Luthra, Aaron J. Ho, Aaron R. Chen, Karthik Guruvayurappan, Michael H. Lorenzini, Carolyn O’Connor, Graham McVicker

## Abstract

The *GATA3* gene is essential for T cell differentiation and is surrounded by risk variants for immune traits. Interpretation of these variants is challenging because the regulatory landscape of *GATA3* is complex with dozens of potential enhancers spread across a large topological associating domain (TAD) and gene expression quantitative trait locus (eQTL) studies provide limited evidence for variant function. Here, we perform a tiling deletion screen in Jurkat T cells to identify 23 candidate regulatory elements. Using small deletions in primary T helper 2 (Th2) cells, we validate the function of five of these elements, two of which contain risk variants for asthma and allergic diseases. We fine-map genome-wide association study (GWAS) signals in a distal regulatory element, 1 Mb downstream, to identify 14 candidate causal variants. Small deletions spanning candidate rs725861 decrease *GATA3* expression in Th2 cells suggesting a causal mechanism for this variant in allergic diseases. Our study demonstrates the power of integrating GWAS signals with deletion mapping and identifies critical regulatory sequences for *GATA3*.

## Main text

T cells orchestrate adaptive immune responses by differentiating into distinct subsets of effector and regulatory T cells. The *GATA3* transcription factor (TF) is central to this process and participates in the differentiation of virtually all T cell subsets. For example, high expression of *GATA3* drives T helper 2 (Th2) cell differentiation^1^, maintains the identity of regulatory T (Treg) cells^2,3^, and disrupts differentiation of T helper 1 and T helper 9 cells^4,5^. *GATA3* has a central role in adaptive immunity and genome-wide association studies (GWAS) have uncovered many nearby genetic variants that are significantly associated with immune-related traits including rheumatoid arthritis^6,7^, multiple sclerosis^8^, type 1 diabetes^9^, asthma and allergic diseases^10,11^. The risk variants for these traits are non-coding and may affect *GATA3* expression, however their interpretation is challenging because we lack a deep understanding of *GATA3’s* cis-regulatory landscape. The high density of GWAS hits near *GATA3* and its cell-type specific regulation, with different expression profiles in different subsets of T cells, make it an excellent model system for interpreting trait-associated human genetic variation. Here, we integrate functional genomic data with CRISPR-mediated genome deletions to identify regulatory elements for *GATA3* and to illuminate the function of genetic variants associated with allergic diseases.

To survey the transcriptional regulation of *GATA3*, we profiled its expression using a published dataset of sorted immune cells^12^. *GATA3* expression is absent in B cells and monocytes, is moderate in naïve CD4^+^ T cells, and is highest in memory Treg cells and Th2 cells, consistent with its established role in Th2 cell differentiation and maintenance^1,13^ (Fig. 1a, b). This motivates our decision to study the regulation of *GATA3* specifically in T cells.

**Fig. 1.**
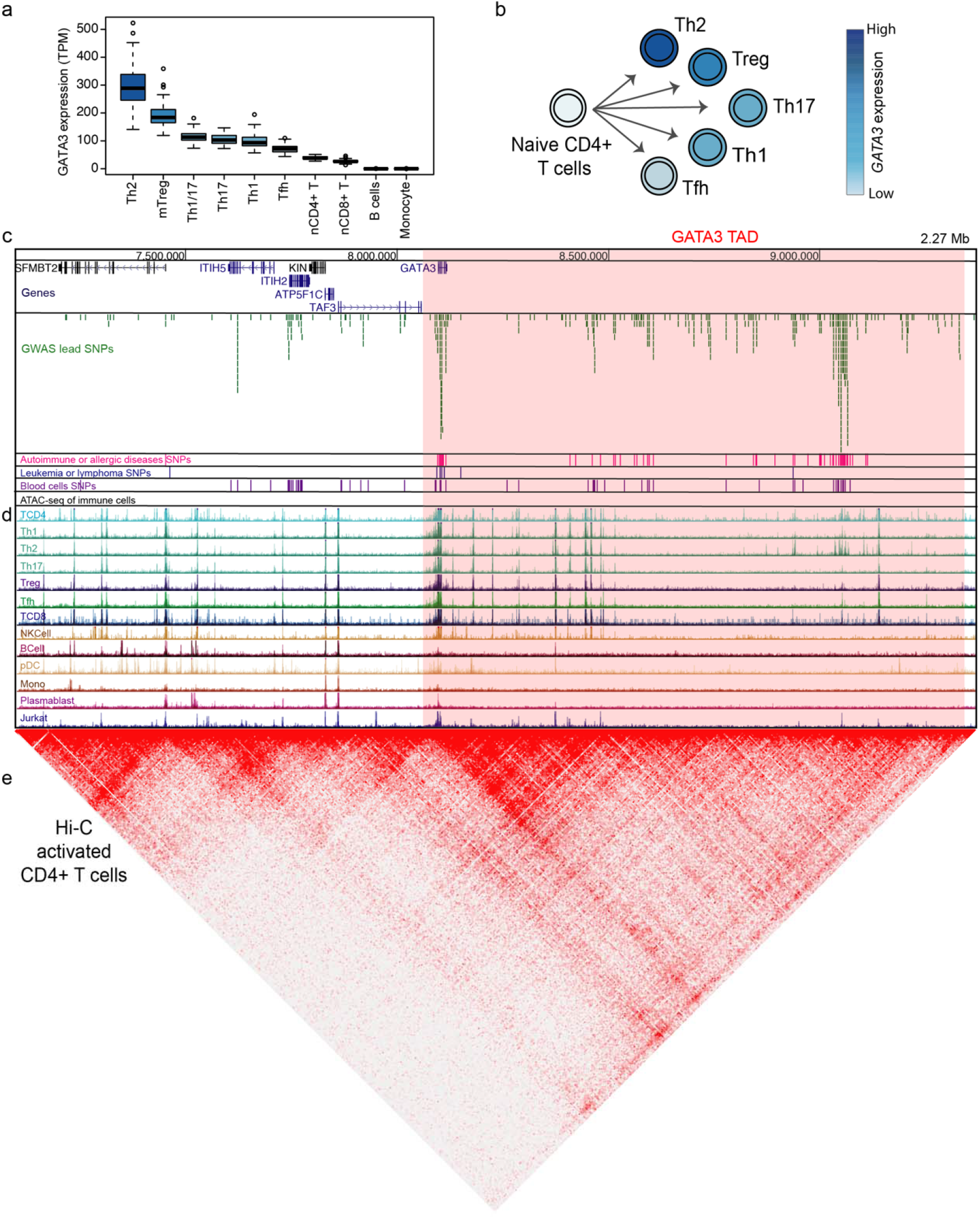
Overview of *GATA3* expression, chromatin accessibility and nearby trait associations. **a**, *GATA3* expression in transcripts-per-million (TPM) in different immune cells from the database of immune cell expression (DICE)^12^. **b**, Differentiation of naïve CD4^+^ T cells into effector and regulatory T cells, with shading indicating *GATA3* expression level. **c**, Lead single-nucleotide polymorphisms (SNPs) from the genome-wide association study (GWAS) catalog in a 2.3 Mb window around *GATA3*. Lead SNPs belonging in three categories (autoimmune or allergic diseases, leukemia or lymphoma, and blood cells) are indicated below. **d**, Chromatin accessibility surrounding *GATA3* in immune cells and Jurkat T cells from published ATAC-seq datasets^16,61^. **e**, Chromatin contact map of the region surrounding *GATA3* from published Hi-C data from CD4^+^ T cells that were activated for 48 hours^15^.

To gain a high-level view of the trait-associated genetic variation surrounding *GATA3*, we examined lead single nucleotide polymorphisms (SNPs) from the GWAS catalog^14^ and classified them into three categories based on immune traits that are potentially mediated by T cells: autoimmune and allergic diseases, leukemia and lymphoma, and blood cell traits (Fig. 1c). The lead SNPs for these trait categories are distributed differently across the 2 Mb region surrounding *GATA3*. Lead SNPs for allergic and autoimmune diseases are located predominantly at *GATA3* and downstream with a major cluster 1 Mb away. Most lead SNPs associated with leukemia and lymphoma are at the *GATA3* gene. Finally, lead SNPs for blood cell traits are scattered across the entire region. The varied locations of lead SNPs for different traits suggest that different regulatory sequences may be involved in the different trait categories.

Given that active regulatory sequences may physically contact their target genes and are typically in open chromatin, we examined published data from Hi-C performed in activated CD4^+^ T cells^15^ and from the assay for transposase accessible chromatin (ATAC-seq) performed in sorted populations of immune cells^16^. These data reveal that *GATA3* is located near the boundary of a large (~1.3 Mb) topological-associating domain (TAD) that extends downstream of the gene (Fig. 1d-e). Dozens of accessible chromatin regions are scattered throughout this TAD and could be considered candidate regulatory elements for *GATA3*. However, this analysis falls short of functionally testing whether these elements regulate *GATA3* expression.

To test if lead SNPs for several immune traits are associated with the expression of *GATA3*, we examined data from gene expression quantitative trait locus (eQTL) studies^12,17,18^. However, the eQTL data provide only weak evidence that GWAS hits affect *GATA3* expression in T cells (Supplementary Table 1). One possible explanation is that power to detect eQTLs is limited by the modest effects of common variants on gene expression and by the relatively small sample sizes of eQTL studies in relevant cell types^19^.

We reasoned that small genomic deletions could determine which sequences regulate *GATA3* expression and could overcome some of the limitations of eQTL studies. Specifically, deletions can directly test the effect of sequences on *GATA3* expression (overcoming ambiguity caused by linkage disequilibrium) and are more likely to have large effects on gene expression than common variants used in eQTL studies. We therefore performed a paired-guide tiling deletion CRISPR/Cas9 screen^20,21^ to discover regulatory sequences that control *GATA3* expression in T cells. With sufficient deletion efficiency, paired-guide screens can cover much larger genome regions than single guide screens or base-editor screens, which have previously been used to screen selected regulatory elements^22–25^. Moreover, unlike regulatory mapping with CRISPR interference or CRISPR activation^26–31^, deletion-based mapping can detect functional sequences that may be insensitive to epigenetic silencing or activation.

To screen for regulatory sequences, we designed 14,769 pairs of single guide RNAs (sgRNAs) to tile across a 2 Mb genome region centered on *GATA3*. The guide pairs target sites separated by a median distance of 1043 bp and are intended to introduce small genome deletions that span the two target sites. The median step size of the deletions is 96 bp, such that each base in the screened region is covered by a median of 8 intended deletions (Supplementary Fig. 1). We note however, that due to variation in guide efficiency and non-homologous end joining (NHEJ), paired guides introduce spanning deletions only some fraction of the time. Instead, they often introduce small insertions/deletions (indels) at each of the target sites^32,33^. To address this, we estimated the deletion efficiency of our system by performing paired-guide mediated deletions of several 1-2 kb sequences in the 2 Mb survey region and measuring the deletion efficiency by quantitative PCR (qPCR) (Supplementary Fig. 2). We estimate the spanning deletion efficiency to be 20-25% and account for this in our analysis described below.

We performed our paired guide screen in Jurkat T cells. While Jurkat is a T cell leukemia cell line, it is a useful model system because the chromatin landscape surrounding *GATA3* mirrors that of primary T cells (Fig. 1d, Supplementary Fig. 3) and it has been previously used to discover disease-relevant T cell regulatory elements^29^. To perform the screen, we cloned oligos encoding the sgRNAs and spCas9, generated a lentiviral library, and transduced the library into Jurkat T cells (Fig. 2a). We selected for cells with viral genome integration and flow sorted them into pools based on GATA3 protein expression (Supplementary Fig. 4). Finally, we performed deep sequencing of the sgRNA pairs in each expression pool (Fig. 2a). We conducted four biological replicates of the screen, sorting into three expression level pools in the first two replicates and seven expression level pools in the other two replicates (Fig. 2b).

**Fig. 2.**
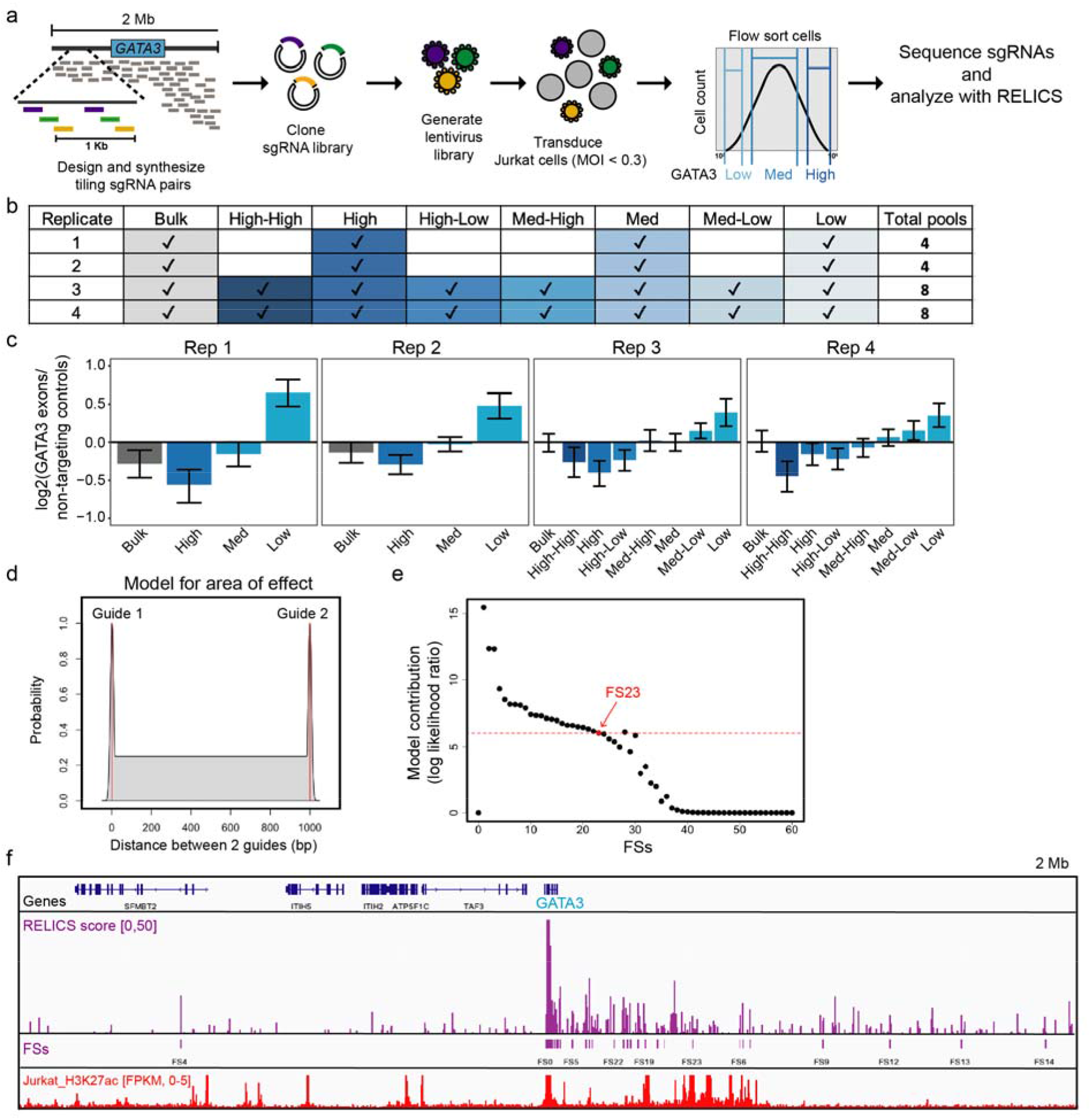
Tiling deletion screen for *GATA3* regulatory elements. **a**, Schematic of CRISPR/Cas9 tiling deletion screen performed in Jurkat T cells. **b**, Four biological replicates of the screen were performed and cells were sorted into three or seven pools based on GATA3 expression. **c**, Guide pairs targeting *GATA3* exons are depleted in low expression pools and enriched in high expression pools compared to nontargeting control guides. Plots for each replicate show the log ratio of the estimated guide pair proportions in each pool. **d**, Area of effect model used by the RELICS analysis, which allows for indels at each guide RNA target site and lower-frequency spanning deletions between target sites. **e**, Improvement in RELICS log likelihood with the inclusion of each functional sequence (FS) and cutoff used to determine significant FSs. **f**, Genome tracks across the 2 Mb screened region showing protein-coding genes, RELICS scores from the tiling deletion screen, FSs predicted by RELICS, and H3K27ac ChIP-seq data from Jurkat cells in fragments per kilobase per million mapped reads (FPKM).

To examine the data generated by the screen, we estimated the proportion of sgRNA pair counts in each pool and compared the sgRNA pairs targeting *GATA3* exons (foreground) to non-targeting control sgRNAs (NTCs) included in the screening library. As expected, sgRNA pairs targeting *GATA3* exons are depleted in the high expression pools and enriched in the low expression pools (Fig. 2c). In contrast, sgRNAs targeting sequences outside of the gene (background), are only slightly enriched in the low expression pool compared to NTCs (Supplementary Fig. 5), which is consistent with a small fraction of them targeting regulatory sequences for the gene. These results indicate that the replicates provide consistent results and that the sgRNA counts can be used to detect sequences that affect *GATA3* expression (Fig. 2c).

To discover regulatory sequences that affect *GATA3* expression, we jointly analyzed the screen data across all of the pools and replicates using RELICS^34^. RELICS is designed to discover functional sequences (FSs) from tiling CRISPR screens and includes features for modeling programmed deletions. RELICS can also leverage data from multiple pools to detect FSs that cause smaller changes in gene expression. RELICS has been extensively validated on experimental data and outperforms other tiling CRISPR screen analysis methods^34^. When running RELICS, we used an area of effect model that assumes a spanning deletion efficiency of 25% to account for the variable mutation events generated by NHEJ (Fig. 2d and Supplementary Fig. 2).

In total, RELICS predicts 23 FSs that affect *GATA3* expression in Jurkat T cells, under a log likelihood ratio threshold of 6 (P=5e-4 by likelihood ratio test) (Fig. 2e-f). These FSs are distributed asymmetrically, with all but one located within the TAD that contains *GATA3*. Most FSs are located within 0.5 Mb of *GATA3*, with some much farther away, and a substantial fraction of them overlap with H3K27ac peaks in Jurkat cells (Fig. 2f).

A limitation of our paired-guide deletion screen is that it was performed in the Jurkat cell line which is less physiologically relevant than primary cells. We therefore view the FSs predicted by RELICS as candidate regulatory sequences and proceeded to validate a subset of the candidates in primary T cells. We selected Th2 cells for validation experiments because *GATA3* expression is highest in Th2 cells (Fig. 1b), and most candidate regulatory sequences (marked by accessible chromatin or H3K27ac) that are present in other T cell subsets are also present in Th2 cells (Figs. 1d, and 3a). We isolated human naïve CD4^+^ T cells, differentiated them into Th2 cells^35^, and electroporated them with pairs of spCas9 ribonucleoproteins (RNPs)^36,37^ (Fig. 3b). An advantage of small deletions is that they can improve the resolution of RELICS’ predictions, which are typically ~1.2kb due to the variable deletion products and efficiencies. As controls, we targeted a “safe harbor” (SH) sequence downstream of *GATA3* with no predicted regulatory function for *GATA3* (negative control) and targeted a *GATA3* exon with a single sgRNA (positive control) (Fig. 3c). We verified the products generated by each deletion experiment using Sanger sequencing and tracking of indels by decomposition (TIDE) (Supplementary Figs. 6 and 7)^38^. Using this system, we generate small (≤ 130bp) deletions with high (40-89%) efficiency (Fig. 3d and Supplementary Fig. 6).

**Fig. 3.**
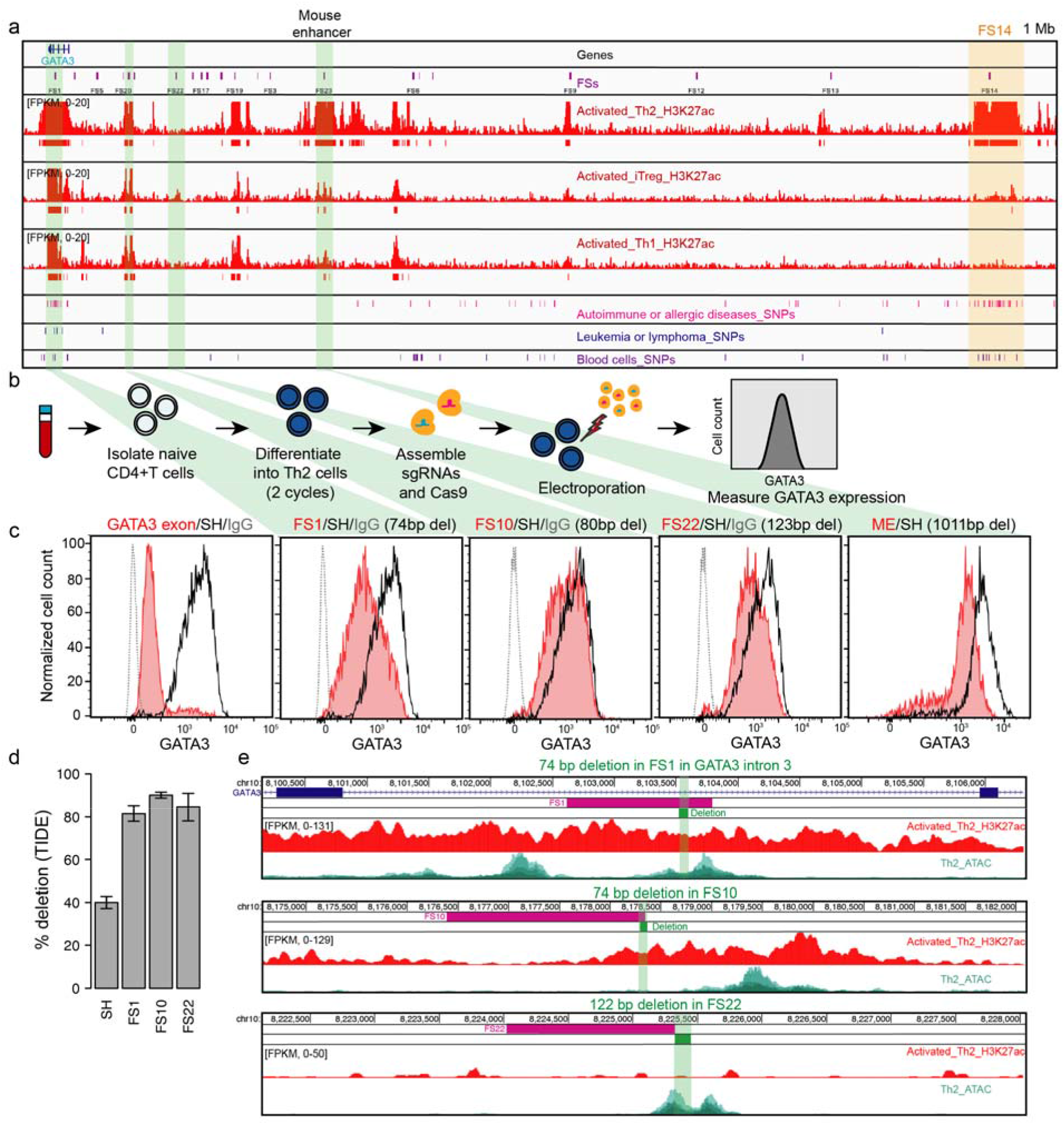
Validation of predicted regulatory elements in primary Th2 cells. **a**, Genome tracks showing the half of the screened region that is downstream of *GATA3*. Tracks are genes, functional sequences (FSs) predicted by RELICS, H3K27ac ChIP-seq for activated Th2 cells, activated induced T regulatory cells (iTregs), and activated Th1 cells from a published study^52^, genome-wide association study (GWAS) lead single nucleotide polymorphisms (SNPs) for different categories of traits. Green highlights indicate regions selected for validation experiments. **b**, Schematic of Cas9 ribonucleoprotein (RNP) validation approach in in vitro differentiated Th2 cells. **c**, Flow cytometry results from intra-cellular staining of GATA3 protein following Cas9 RNP experiments. Each panel also shows results using an isotype control (IgG, dotted line) and from targeting a safe harbor (SH) genome sequence (black outline). Results from independent replicate experiments are provided in Supplementary Fig. 8. **d**, Estimated percentage of cells carrying targeted deletions for SH, FS1, FS10, and FS22 (quantified by Tracking of Indels by Decomposition (TIDE)^38,62^. **e**, Zoom-in tracks showing the functional sequences targeted for deletions. Genome tracks showing *GATA3* gene, FSs, H3K27ac ChIP-seq for activated Th2 cells, and ATAC-seq of Th2 cells (overlay with results from multiple donors).

We first targeted the top-ranked prediction, FS1, for validation in Th2 cells. FS1 is located within the third intron of *GATA3* and overlaps a strong H3K27ac peak. We designed a pair of sgRNAs to delete a 74 bp sequence from FS1, which contains an ATAC-seq peak in Th2 cells (Fig 3e). Deletion of this sequence reduces GATA3 expression substantially (Fig. 3c and Supplementary Figs. 7-8) indicating that it acts as an enhancer for *GATA3*.

FS23 is orthologous to a mouse genome sequence that was previously identified as a strong enhancer for *GATA3* in mouse T cells^39^. In addition, this sequence overlaps with ATAC-seq peaks in all T cell subsets and has strong H3K27ac signals in Jurkat cells, naïve CD4^+^ T cells, and Th2 cells. To test the function of this ‘mouse enhancer’ sequence, we performed spCas9 RNP experiments to delete a 1011 bp of the sequence. We found that GATA3 expression is reduced compared to SH control experiments, confirming that this sequence acts as an enhancer in human T cells (Fig. 3c).

Finally, we deleted 80 bp of FS10 and 123 bp of FS22. Both deletions decrease *GATA3* expression (Fig. 3c and Supplementary Fig. 8). FS10 is marked by H3K27ac and acts as another classical enhancer in Th2 cells (Fig 3e). FS22 lacks H3K27ac signals, but it does overlap with an ATAC-seq peak (Fig 3e), suggesting that it may function differently from a canonical enhancer.

To determine whether regulatory elements identified in our screen can help interpret GWAS variants, we intersected the FSs with GWAS hits and observed that FS1 and FS14 are located within two distinct clusters of risk variants associated with autoimmune and allergic diseases (Fig. 3a). We examined the region surrounding FS14, which is almost 1 Mb downstream of *GATA3*, and which has a high density of lead SNPs (Fig. 3a). This region is contained within a broad 44 kb H2K27ac domain, which is present in Th2 cells but not in other T cell subsets, suggesting that it may be a distal Th2-specific enhancer (Fig. 3a). Because Th2 cells have an established role in allergic diseases^40^ and high *GATA3* expression is required for differentiation and maintenance of Th2 cells^1^, we decided to (i) analyze recent GWASs for asthma and allergic diseases (allergic rhinitis or eczema)^10^, and (ii) test the function of sequences containing risk variants in in vitro differentiated Th2 cells.

A single GWAS hit for asthma is situated ~1 Mb downstream of *GATA3* (Fig. 4a) and a pair of independent GWAS hits for allergic diseases are located at ~400 kb and ~1 Mb downstream of *GATA3*. We name these hits risk region 1 and risk region 2 (Fig. 4a,b) and focus on risk region 1, because it is associated with both traits and overlaps the large Th2-specific enhancer-like sequence described above. We performed fine-mapping to identify 95% credible sets (CSs) containing candidate causal variants using SuSiE^41^. For asthma, we identified two CSs: CS1, which contains three candidate causal SNPs, and CS2, which contains 11 candidate SNPs (Fig. 4c). For allergic diseases, we identified a single CS containing the same three candidate SNPs as CS1 (Fig. 4d, 5a).

**Fig. 4.**
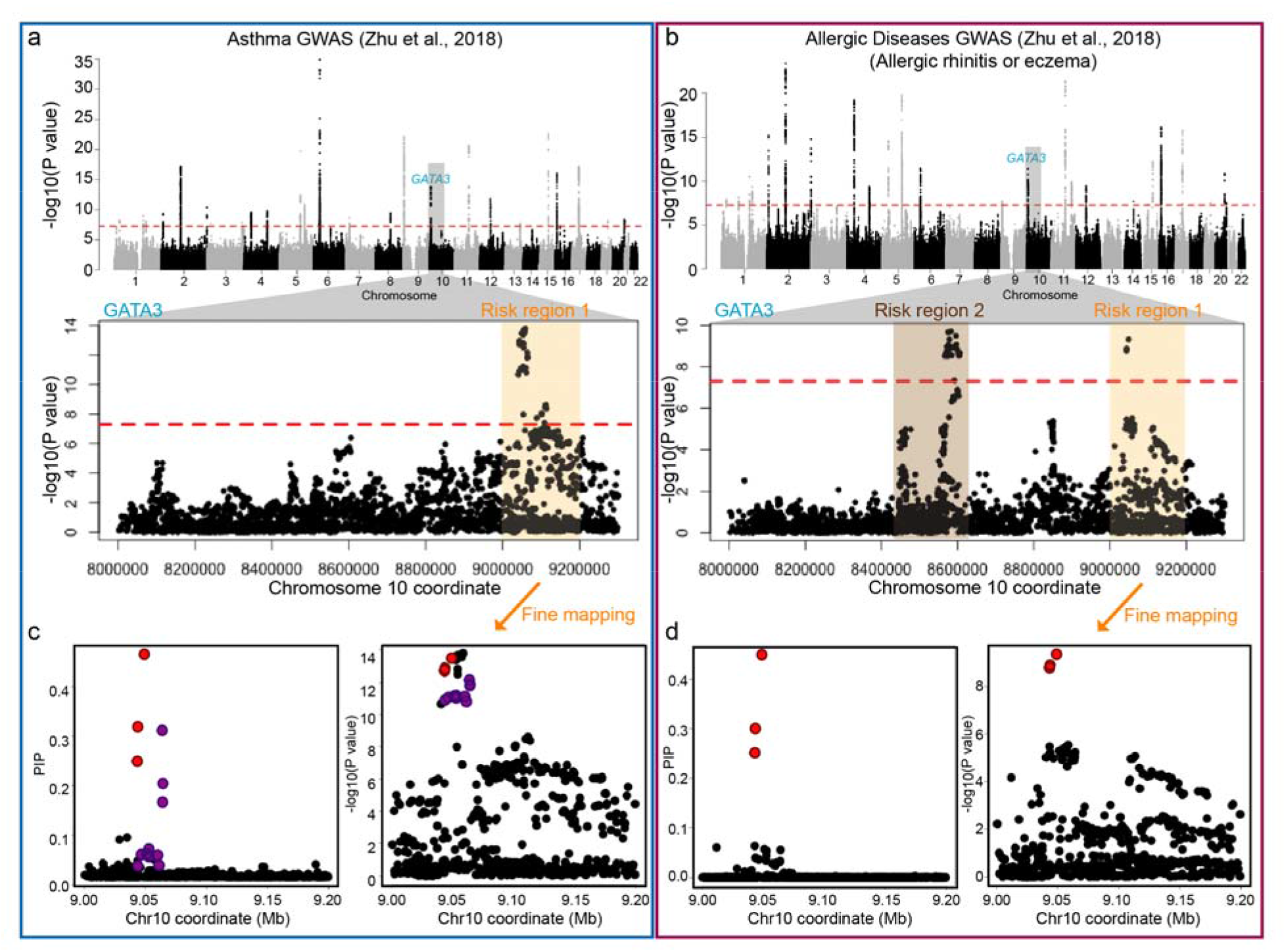
Fine-mapping of genome-wide association study hits at the *GATA3* locus. **a**, Manhattan plot for a published genome-wide association study (GWAS) for asthma^10^, and zoom-in of the *GATA3* locus showing a single region of trait association “risk region 1” located ~1 Mb downstream of the gene. **b**, Manhattan plot and zoom-in for a published GWAS of allergic diseases^10^ showing two regions of trait association downstream of *GATA3*, “risk region 1” and “risk region 2”. **c**, Fine-mapping of risk region 1 with SuSIE^41^ reveals two credible sets (CSs) for asthma association highlighted in red and purple. Left panel: posterior inclusion probabilities (PIP) for variants in risk region 1. Right panel: −log10 p-values of the same variants. **d**, Fine-mapping of allergic disease reveals a single credible set, highlighted in red, containing the same variants as CS1 for asthma.

**Fig. 5.**
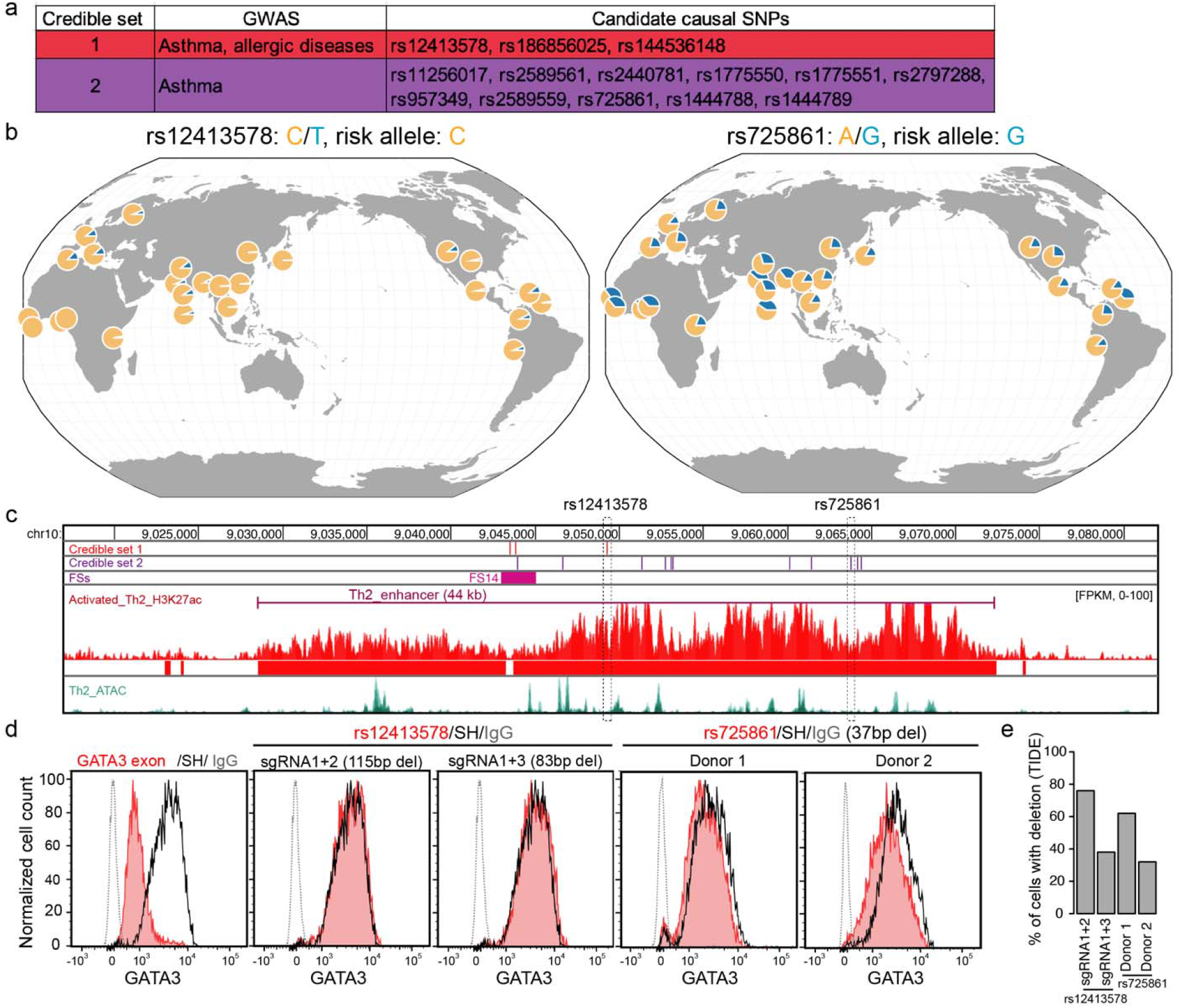
Candidate variants for allergic diseases and asthma in a distal Th2 enhancer. **a**, Candidate causal single nucleotide polymorphisms for asthma and allergic diseases that make up credible sets (CSs) CS1 and CS2. **b**, Worldwide population allele frequencies for the SNPs with the highest posterior inclusion probability in each CS. Allele frequency estimates are from the 1000 genomes project^63^ and plotted with the Geography of Genetic Variants Browser^64^. **c**, Genome tracks showing the locations of the candidate variants, predicted functional sequences (FSs), H3K27ac ChIP-seq for activated Th2 cells, and ATAC-seq for Th2 cells. The 44 kb distal Th2 enhancer is indicated. **d**, Flow cytometry results following electroporation of Cas9 ribonucleoproteins (RNPs) targeting an exon of *GATA3* or small deletions spanning candidate SNPs rs12413578 or rs725861 in in vitro differentiated Th2 cells. Two pairs of gRNAs that differ in one gRNA were used to delete 115 bp or 83 bp spanning rs12413578. A 37 bp sequence spanning rs725861 was deleted in Th2 cells from two different donors. **e**, Estimated percentage of cells carrying targeted deletions for samples in **d** (quantified by Tracking of Indels by Decomposition IDE^38^ using the Synthego ICE analysis tool^62^).

We examined the worldwide allele frequencies of the SNPs with the highest posterior inclusion probabilities (PIP) in CS1 (rs12413578) and CS2 (rs725861). For rs12413578, the risk allele is the major allele, and the protective allele has the highest allele frequencies in populations with European ancestry and the Indian subcontinent (Fig. 5b). In contrast, the risk allele for rs725861 is the minor allele and has the highest frequencies in Western Africa and the Indian subcontinent (Fig. 5b).

To test the function of candidate SNPs in the two CSs, we designed pairs of guides to delete small sequences surrounding 6 of the 14 candidates. Only some of these pairs of guides yielded high-efficiency deletions, but fortunately these deletions encompassed the SNPs with the highest PIP in each CS (Fig 5c). We deleted 115 bp and 83 bp sequences surrounding rs12413578, the highest PIP SNP in CS1, in in vitro differentiated Th2 cells. Neither of these deletions affected GATA3 expression in Th2 cells (Fig. 5d-e and Supplementary Fig. 9), suggesting that the sequence containing rs12413578 may not regulate *GATA3* expression in the Th2 condition we tested. In contrast, deletion of a 37 bp sequence surrounding rs725861, the highest PIP SNP in CS2, decreased GATA3 expression in Th2 cells from two different donors (Fig. 5d-e and Supplementary Fig. 9). This indicates that this distal sequence functions as an enhancer in Th2 cells and that risk variants for allergic diseases and asthma are likely to affect *GATA3* expression in this cell type.

In response to pathogens, naïve CD4^+^ T cells become activated and differentiate into effector and regulatory T cell types to mount appropriate immune responses. However, incorrect immune responses lead to autoimmune and allergic diseases. Our results show that expression of *GATA3*, a key regulator of T cell differentiation, is coordinated by a large pool of downstream regulatory sequences and demonstrate the power of high-throughput deletions coupled with computational analysis and validation experiments in primary cells. Sequences detected by this approach can be remarkably useful for the interpretation of trait-associated genetic variation, and we provide evidence that risk variants for allergic diseases affect the function of a Th2-specific *GATA3* enhancer.

## Methods

### Antibodies

All antibodies used in the study for fluorescence activated cell sorting, flow cytometry, and cell culture are listed in Supplementary Table 2.

### Oligos

All oligos used in this study are listed in Supplementary Table 3.

### Cell culture

All cells were cultured at 37°C in a humidified incubator with 5% CO_2_. Jurkat cells (Clone E6-1) were a gift from Bjorn Lillemeier (Salk Institute for Biological Studies, La Jolla, CA). Primary human T cells were sourced from fresh buffy coats from two de-identified blood donors collected by the San Diego Blood Bank (San Diego, CA). Jurkat cells were cultured in RPMI-1640 medium (Gibco) supplemented with 5% fetal bovine serum (FBS, Gemini). Peripheral blood mononuclear cells (PBMCs) were isolated from the buffy coats using Ficoll (Lymphoprep solution, Cosmo Bio #AXS-1114545) density gradient centrifugation. Naïve CD4^+^ T cells were isolated from frozen PBMCs using Naïve CD4^+^ T cell isolation kit II, human (Miltenyi Biotec #130-094-131). Unstimulated naïve CD4^+^ T cells were cultured in IMDM (Gibco) with 5% heat-inactivated FBS, 2% human AB serum (Gemini #100-512), and 40 U/ml recombinant human IL-2 (Miltenyi Biotec, #130-097-745). HEK293FT cells were a gift from the Salk viral vector core and were cultured in DMEM (high glucose, Life Technologies #10569010) containing 10% FBS, 0.1 mM MEM™ non-essential amino acid (NEAA), 6 mM L-glutamine, 1mM MEM™ sodium pyruvate, and 500 mg/ml Geneticin (Life Technologies #10131035).

### T helper 2 cell differentiation

In vitro T helper 2 (Th2) cell differentiation was carried out following the protocol published in Schmiedel et al. 2016^35^. Naïve CD4^+^ T cells were cultured with Human T-Activator CD3/CD28 Dynabeads (Invitrogen #11161D) at a bead-to-cell ratio of 1:1 in the presence of recombinant IL-4 (Fisher Scientific, #204-IL-010, 10 ng/ml) and anti-IFN-γ antibody (10 μg/ml). The anti-CD3/CD28 Dynabeads were removed after 48 hours of stimulation and the cells were expanded in culture with 40 U/ml recombinant human IL-2 for 3 more days before introducing dual spCas9/sgRNA ribonucleoprotein (RNP) complexes.

### Guide pair design for the tiling deletion screen

To design guide sequences, we extracted a 2.02 million base pair input region from human genome assembly hg19 (*GATA3* gene region plus 1MB flanking on either side chr10: 7,096,667-9,117,164). Using this sequence, we extracted a list of all possible candidate guides of length 19-21bp that were adjacent to a spCas9 protospacer-adjacent motif (PAM) (NGG, for forward strand guides and CCN for reverse strand guides). We reduced the set of candidate guides to those with a G in first position, because a leading G provides the most efficient expression from a U6 promoter.

Once a list of candidate guide sequences was established, we computed specificity scores for each guide so that off-target effects could be reduced. We aligned guides to the genome with BWA^42^, allowing up to three mismatches to the reference sequence. Using the genome alignments, we calculated a specificity score for each candidate guide using the method described by Hsu et al. 2013^43^. We then discarded all guides with a specificity score less than 20, which retained enough guides to select reasonably-spaced guide pairs, while reducing off-target effects.

To choose pairs of guides for tiling deletions we defined a minimum step size (S; distance from first target site from previous pair of guides) and deletion size (D; distance between two target sites). When designing each deletion, we chose the pair of guides from our filtered guide list such that the step size and deletion size thresholds were exceeded by the smallest possible amount. We set S to 65 bp and D to 1000 bp, which gave us a total of 14,555 targeting guide pairs with a median size of 1043 bp for D and median step size of 96 bp for S, such that each base in the screened region would be covered by an average of 8 programmed deletions (Supplementary Fig. 1). In addition, we designed 214 non-targeting negative control guide pairs, which were random sequences that did not align to any sequence in the reference human genome.

### CRISPR/Cas9 tiling deletion screen

Our screening method was based on the published CREST-seq method^20^, which we optimized for Jurkat cells with minor modifications. Briefly, an oligo pool encoding 14,769 guide pairs was synthesized by Agilent Technologies. We PCR amplified the oligo pool for 25 cycles using KAPA HiFi DNA polymerase (Roche, KK2103). We digested lentiCRISPRv2 plasmid (Addgene #52961) with BsmBI (Thermo Scientific, #FERFD0454) and cloned the purified oligo pool to linearized lentiCRISPRv2 plasmid by Gibson assembly for 1 hour incubation at 50 °C. The end product was precipitated by ethanol and quantified by Qubit 3 Fluorometer (Thermo Fisher Scientific). The product was electrotransformed into Endura™ Competent Cells (Lucigen #60242) and grown on big agar plates. 1.51 million bacterial colonies were collected and the plasmids were extracted with NucleoBond Xtra Maxi EF kit (Takara Bio USA #740424). To insert a TmU DNA fragment containing the tracRNA (E/F) and the mouse U6 promoter (mU6) sequences (synthesized by IDT gBlock service) into the plasmid library, the plasmid library was linearized by BsmbI digestion and ligated with TmU fragments with Quick ligation™ kit (New England BioLabs, M2200S) for 5 minutes at room temperature. The final plasmid library was electrotransformed into Endura™ Competent Cells (Lucigen #60242) and grown on big agar plates. More than 1.5 million bacterial colonies were collected and the plasmids were extracted with NucleoBond Xtra Maxi EF kit. The pairs of guides in the final plasmid library were deep sequenced on an Illumina NextSeq instrument to confirm the final plasmid library preserved the pairing and frequency of the guide library.

To generate lentiviral libraries, HEK293FT cells were seeded at about 90% confluence in 10-12 10-cm dishes. Two hours before transfection, the media of the HEK293FT cells was replaced with the Jurkat cell media (RPMI-1640 containing 5% FBS). For each 10 cm dish, 12 μg of plasmid library was cotransfected with 9 μg psPAX2 and 3 μg pMD2.G (Addgene #12260 and #12259) into HEK293FT cells using 72 μl Lipofectamine 2000 (Life Technologies #11668019). Transfectants were removed after 16 hours of incubation and replaced with Jurkat cell media, 5ml/dish. The virus-containing media was harvested twice at 48 and 72 hours after transfection. The two harvests were combined and centrifugated at 1200 rpm for 5 minutes. The virus-containing supernatants were collected and small aliquots of viruses were stored at −80°C.

To titrate the viral solution, 0.1 million Jurkat cells were seeded in 0.2 ml media per well in a 96-well plate. Different volumes (0, 2, 4, 8, 16 and 32 μl) of viral solution were added to each well with two replicates to infect Jurkat cells by spinoculation. 72 hours after transduction, Jurkat cells were treated with 0.8 μg/ml puromycin (Life Technologies #A1113802) for 48 hours. The live cell number of each well was counted to calculate the viral titer.

We carried out four independent replicate screens. In each screen, 50 million Jurkat cells were transduced with lentivirus library at a low multiplicity of infection (MOI < 0.3) to ensure that most cells received at most one viral particle (one pair of guides). Jurkat cells were spinoculated in 50 ml conical tube with the lentivirus library at 800×g for 90 minutes at 32°C with 0.8 μg/ml polybrene (EMD Millipore #TR-1003-G) and 10 μM HEPES (Life Technologies #15630080). Each 50 ml conical tube used for spinoculation contained 5 million cells in 5 ml media. At 24 hours post-transduction, the media of the infected cells were replaced with fresh media, and at 72 hours post-transduction, Jurkat cells were treated with 0.8 μg/ml puromycin for 48 hours to select for cells with viral genome integration. Over 90% of cells died from puromycin selection and we observed that the presence of dead cells affected the expression of spCas9 from live cells. To remove the dead cells from the culture, cells were filled up to 10 ml with media in 15 ml canonical tubes. 2ml Ficoll-Paque PLUS Media (GE Healthcare #17144002) was slowly added to the bottom of the tube to create a gradient. Cells were centrifugated at 2000 rpm for 10 minutes at room temperature with acceleration speed of 4 and deceleration speed of 4. A white layer of live cells above the gradient was collected and live cells were centrifugated at 1200 rpm for 5 minutes at room temperature to remove cell debris. The remaining live cells were cultured in conditional media (50% fresh media plus 50% filtered old media) for two days for recovery.

Fluorescence Activated Cell Sorting (FACS) was used to obtain pure populations of infected cells (producing DYKDDDDK tagged spCas9 protein) separated into pools with different GATA3 protein levels. 150-300 Million cells were collected and stained with Zombie Violet™ fixable viability kit (Biolegend #423113) with 1:200 dilution in phosphate-buffered saline (PBS) for 15 minutes at room temperature. For immuno-labeling, True-Nuclear Transcription factor Buffer set (Biolegend #424401) was used according to manufacturer’s recommendations: intracellular labeling with APC anti-GATA3 antibody (2.5 ul antibody per million cells in 100 ul) (Biolegend #653806) and PE anti-DYKDDDDK Tag antibody (0.6 ul antibody per million cells in 100 ul) (Biolegend #637310) for DYKDDDDK-tagged Cas9 was carried out at room temperature for 45 minutes. After washing cells in FACS buffer (1% BSA in PBS), cells were resuspended in FACS buffer and kept on ice where they were taken to the Flow Cytometry Core. For FACS, the following gating strategy was applied: Cells were first gated on Violet™ signal intensity to obtain the live cell population. Live cells were selected on forward scatter (FSC) and side scatter (SSC), then aggregates were excluded using pulse width for FSC and SSC before gating on PE labeling (for Cas9 protein) and lastly, APC signal intensity (GATA3 protein level). Cells were sorted into low, medium, high expression pools of GATA3 for the first 2 replicate screens, and into low, medium-low, medium, medium-high, high-low, high, high-high expression pools of GATA3 for replicate screens 3 and 4 using a BD Influx™ cell sorter (100um nozzle, 17.5PSI using 1 x PBS as sheath fluid). A representative gating strategy for cell sorting based on APC-GATA3 level is shown in fig. S7. A minimum of one million cells were collected for each pool in each replicate in 100 ul PBS.

To extract genomic DNAs, 0.5 million cells were incubated with 20 μl, 10 mg/ml proteinase K (SIGMA-ALDRICH INC, P8044) in 300 μl cell swelling buffer (10mM Tris-HCl, pH 8.0, 85mM KCl, 0.5% NP-40, and 10mM MgCl_2_) at 55 °C overnight. Genomic DNA was extracted using Phenol:Chloroform:Isoamyl Alcohol (Life Technologies #15593031) and ethanol precipitation. The gRNA inserts were amplified from genomic DNA from multiple PCR reactions, in which 400 ng gRNA DNA was used in one single PCR reaction. For example, if there were a total 12 μg genomic DNA in one pool, 30 PCR reactions were performed using Herculase II fusion DNA polymerase (Agilent Technologies, #600679) for 3 cycles to amplify guide insert and add Illumina TruSeq sequencing adaptors. For each pool, the amplified guide inserts from multiple PCR reactions were pooled together and purified by AMPure XP beads (Beckman Coulter). A second PCR was performed to add TruSeq sequencing primers to the purified guide inserts for 16 cycles, with 400 ng/PCR reaction. The final guide libraries were sequenced using an Illumina NextSeq instrument in pair-end mode with 75 bp read length as described in the CREST-seq method^20^.

### Tabulating guide pair counts

We discarded all guides that overlapped with deletions, insertions, translocations and inversions identified by whole-genome sequencing of the Jurkat cell line^44^. In total we excluded 1302 guide pairs by these criteria leaving 13,253 targeting guide pairs for analysis.

To assign sequencing reads to guide pairs and count the number of guide occurrences in each pool, we mapped the reads to the guide library using BWA-MEM^45^. We first created a custom ‘reference genome’ library that contained the complete set of designed guide pairs. Each guide pair in the reference library included flanking 5’ and 3’ sequences from the viral vector. After aligning guides to the custom reference, we retained only those guide pairs with at least 60 matches (as defined by the CIGAR string). Aligned reads where the guide pairs were observed to be recombined^46^ were discarded.

### RELICS analysis of guide pair counts

We used RELICS (https://github.com/patfiaux/RELICS)^34^ to jointly analyze the guide pair counts across all pools and replicates (24 pools total).

RELICS leverages a set of known functional sequences, labeled FS0, to estimate sorting parameters, which describe the probability that cells containing different guides will be sorted into each pool. We labeled the sequences corresponding to *GATA3* exons as FS0. Based on the definition of FS0, RELICS divides all guide pairs into two sets: those predicted to generate deletions that overlap FS0 (foreground), and all other targeting guide pairs (background). Using RELICS, we obtained maximum likelihood estimates of sorting parameters for foreground, background and non-targeting control guides for each pool in each replicate. We plot the log2 ratio of foreground/non-targeting control sorting probabilities in Figure 2c and the log ratio of background/non-targeting control sorting probabilities in Supplementary Figure 5. Confidence intervals for the log ratios were estimated as the central 95% of values from 1000 bootstrap iterations, in which the guide pairs in the foreground, background, and non-targeting control sets were re-sampled with replacement.

RELICS requires that the user specify a maximum number of FSs, and we set this value to 60.

RELICS models the dispersion of observed guide counts as a function of the total count of a guide across pools using fitted splines, with a user-specified number of degrees of freedom. We set the the degrees of freedom to 2, which yielded a good spline fit for all replicates (Supplementary Fig. 10).

For paired guide deletion screens, RELICS allows for the possibility that small insertions/deletions (indels) at each target site may occur instead of the intended deletion between the target sites. In addition, RELICS allows the indels to occur with a probability that varies with distance from the target sites. We specified that the large deletions occur with relatively low efficiency (probability 25%), and an area of effect for indels that followed a normal distribution with standard deviation of 8.5 bp. These parameters were set based on published experimental data regarding the mutational events that result from the use of spCas9 and paired-guides^32,47,48^.

### Peak calling for ChIP-seq and ATAC datasets

For Jurkat cells, ChIP-seq data for H3K27ac was obtained from Hnisz et al. 2016^49^. To process the sequencing data, the adaptors of the reads were trimmed and the quality of the reads were accessed using trim_galore (v 0.6.6), a wrapper for cutadapt (v 1.18)^50^ and fastqc (0.11.9) (https://www.bioinformatics.babraham.ac.uk/projects/fastqc/). Reads were aligned using BWA (v 0.7.17)^42^ using with the bwa mem command and default arguments. The alignments were sorted and alignments with MAPQ score less than 30 were removed using samtools (v 1.12)^51^. Duplicate reads were removed using picard (v 2.25.4) (https://broadinstitute.github.io/picard/).

For in vitro differentiated Th2 and iTreg cells, ChIP-seq data for H3K27ac were obtained from Soskic et al. 2019^52^. We downloaded the alignment data in CRAM format, which we converted to BAM. We then merged BAM files from three donors based on cell type, activation state, and stimulation time before data processing.

Peaks were called using MACS2 (v 2.2.7.1)^53^ with the MACS2 callpeak command. H3K27ac ChIP-seq peaks were called with the --*q 0.01* option, and default arguments. Tracks were created with deepTools (v 3.5.1)^54^.

### CRISPR/Cas9-mediated deletion with synthetic sgRNAs

Guides for individual deletion experiments with spCas9 RNPs were designed using FlashFry^55^, CRISPR-SE^56^, and GuideScan^57^. Single guide RNAs (sgRNAs) with predicted high specificity scores and low off-targeting were selected to generate small deletions (<130bp) at target regions. sgRNAs were synthesized by IDT with modifications on the 5’ and 3’ends to enhance efficiency^37^. For example, for an sgRNA oligo with target sequence GCACTAAATCATTCACTGGG, the final synthesized sgRNA is mG*mC*mA*rCrUrArArArUrCrArUrUrCrArCrUrGrGrGrGrUrUrUrArArGrArGrCrUrArUrGrCrUrGrG rArArArCrArGrCrArUrArGrCrArArGrUrUrUrArArArUrArArGrGrCrUrArGrUrCrCrGrUrUrArUrCrAr ArCrUrUrGrArArArArArGrUrGrGrCrArCrCrGrArGrUrCrGrGrUmG*mC*mU*rU (“r” for a RNA base, “m” for a 2’OMe modified RNA base, “*” for a phosphorothioate bond.). The sequences of the guides targeting safe harbor regions, *GATA3* exon, FS1, FS10, FS22, mouse enhancer which contains FS23, rs12413578 and rs725861 are listed in the Supplemental Table 3. Synthetic sgRNAs were reconstituted in nuclease-free water and stored at −80°C with small aliquots.

To assemble the dual spCas9 RNP complexes, 90 pmole of each sgRNA was mixed with 5 μg SpyFi™ Cas9 Nuclease (Aldevron LLC #9214-0.25MG) or Alt-R® S.p. Cas9 Nuclease V3 (IDT), and incubated for 15 minutes at room temperature. The Cas9 RNPs from the two paired guides were added to one million in vitro differentiated Th2 cells for 5 days in 20 ul MaxCyte® electroporation buffer. The mixture was immediately electroporated using a MaxCyte ATx® instrument with expanded T cell protocol 2. Cells were transferred to fresh media immediately after electroporation. Cells were cultured in IMDM (Gibco) with 5% heat-inactivated FBS, 2% human AB serum (Gemini #100-512), and 40 U/ml recombinant human IL-2 (Miltenyi Biotec, #130-097-745) for 2 days prior to flow cytometry analysis.

### CRISPR/Cas9-mediated deletion efficiency estimation by qPCR

Approximately 0.5 million cells were collected and incubated with 20 μl, 10 mg/ml proteinase K (Sigma-Aldrich Inc, P8044) in 300 μl cell swelling buffer (10mM Tris-HCl, pH 8.0, 85mM KCl, 0.5% NP-40, and 10mM MgCl_2_) at 55 °C overnight. Genomic DNA was extracted by Phenol:Chloroform:Isoamyl Alcohol (Life Technologies #15593031) and precipitated by ethanol precipitation. To quantify the deletion efficiency, qPCR was performed to quantify the relative ratio of a genomic region with targeted deletions in samples compared to the control without deletions. Each deletion was quantified with 4-6 qPCR replicates. For each sample, 2.5-10 ng of genomic DNA was used in a single 10 or 20 μl qPCR reaction using PowerUP™ SYBR™ green master mix (Life Technologies #A25742) and analyzed by a QuantStudio 6 Flex real-time PCR system (Thermo Fisher Diagnostics).

### Flow cytometry analysis

CRISPR/Cas9 RNP-treated Th2 cells were stained intracellularly with APC anti-GATA3 antibody (Biolegend #653806, 1:40 dilution) at room temperature for 30 minutes using True-Nuclear Transcription Factor Buffer Set (Biolegend #424401) to assess GATA3 protein level. The GATA3 protein levels for each cell were assessed with a BD FACSCanto II™ cytometer. A representative gating strategy for flow cytometry analysis is shown in Supplementary Fig. 4.

### Tracking of Indels by Decomposition (TIDE)

TIDE experiments were carried out based on the method described in Brinkman et al., 2014^38^. Genomic DNAs from CRISPR edited cells were extracted using Quick-DNA Microprep kit (Zymo Research, D3020). PCR primers were designed to amplify ~1000 bp amplicons from 100-200bp upstream of targeted deletions to 800-900bp downstream of target deletions. PCR reactions were performed using KAPA HiFi kits (Roche Sequencing) with 40-100 ng genomic DNA and GC buffer for 25 cycles. PCR products were purified using AMPure beads (Beckman Coulter) and sent for Sanger sequencing by Azenta Life Sciences. The sequencing results were analyzed using the ICE CRISPR Analysis Tool (Synthego)^58^.

### Fine-mapping

Summary statistics from GWAS for allergic diseases were obtained from Zhu et al. 2018^10^. The linkage disequilibrium (LD) correlation matrix was computed using a sample of 10,000 individuals from the UK Biobank. Statistical fine-mapping was performed separately on the two risk regions downstream of *GATA3* using SuSiE (https://github.com/stephenslab/susieR)^41^ using the summary statistics and LD correlation matrix. Fine-mapping of the 200 kb region surrounding each risk region (hg19 chr10: 9,000,000 - 9,200,000 and chr10: 8,440,000 - 8,640,000) was performed using the Susie.rss function, allowing for a maximum of 15 causal variants and a maximum of 2000 iterations to yield 95% credible sets.

### GWAS eQTL intersection analysis

Summary statistics for genome-wide significant (P < 5e-8) lead SNPs within 1 MB of *GATA3* were downloaded from the GWAS catalog^14^. We selected a subset of studies to represent a spectrum of immune disease traits (multiple sclerosis^8^, rheumatoid arthritis^59^, type 1 diabetes^9^, allergic disease and asthma^10^), choosing the GWAS with the largest sample size when there were multiple hits for the same trait. We then intersected the lead SNPs from these studies with uniformly processed immune cell eQTL data from BLUEPRINT^60^ and DICE^12^ downloaded from the eQTL catalog^18^. We report P-values for ‘gene expression’ eQTL associations for each GWAS lead SNP in Supplementary Table S1.

## Acknowledgments

We thank B. Ren and Y. Diao for assistance with the planning of our tiling deletion screen, Y. Zheng and Z. Liu for helpful discussions, P. Hsu and S. Konermann for advice about CRISPR screens, MaxCyte for their assistance with electroporation, and N. Hah and the Salk Next Generation Sequencing Core for their technical support with sequencing. This study was supported by NIH/NHGRI grant HG011315 to GM; NIH/NIDDK grant DK122607; NIH/NIAID grant AI107027; the National Cancer Institute funded Salk Institute Cancer Center (NIH/NCI CCSG: 2 P30 014195); the 2020 Salk Women & Science Award to HVC; the 2020 Salk Alumni Fellowship Award to HVC; the H.A. and Mary K. Chapman Charitable Trust Fellowship to PCF; the Jesse and Caryl Philips Foundation Fellowship to PCF; the Pioneer Fund Postdoctoral Scholar Award to AS; and the Frederick B. Rentschler Developmental Chair to GM. Sequencing and flow cytometry were carried out by the NGS and Flow Cytometry Core Facilities of the Salk Institute with funding from NIH/NCI CCSG: P30 014195, the Chapman Foundation, and the Helmsley Charitable Trust.

## Author contributions

HVC and GM conceived of the project. HVC performed the tiling deletion screen and all other experiments, drafted the initial version of the manuscript, analyzed flow cytometry and GWAS data, and made the figures. GM supervised the project, acquired funding for the project, and edited and wrote the manuscript with HVC. ARC assisted with experiments under the direction of HVC. AJH performed computational analyses of functional sequences and bioinformatics processing of ChIP-seq and ATAC-seq data. PCF and KG applied and updated RELICS to analyze the screening data and generated some of the screening figures. AS performed analysis of ATAC-seq data and sequence data and provided comments on the manuscript. MHL assisted with electroporation, flow cytometry and TIDE experiments, provided input on T cell editing, and provided comments on the manuscript. IL wrote scripts to design the guide RNAs used in the screen. CC assisted with flow cytometry experiments.

## Competing interests

Electroporation experiments were performed with a MaxCyte ATx instrument that was provided to the McVicker laboratory by MaxCyte for technology development and evaluation purposes. All authors declare that they have no other competing interests.

## Data and materials availability

All data are available in the main text or the supplementary materials. The raw sequencing data for the tiling deletion screen have been deposited in GEO under accession GSE190860.

## Notes

https://www.ncbi.nlm.nih.gov/geo/query/acc.cgi?acc=GSE190860

