## Supplementary Materials for "Deletion mapping of regulatory elements for *GATA3* reveals a distal T helper 2 cell enhancer involved in allergic diseases"

**This PDF file includes:**

Supplementary Figs. 1 to 10
Supplementary Table 2. List of antibodies used in this study

**The following Supplementary Material is provided as separate files:**

Supplementary Table 1. A table for gene expression quantitative trait locus (eQTL) studies
Supplementary Table 3. List of oligos used in the study
Supplementary Table 4. List of guide pairs used in the GATA3 tiling deletion screen
Supplementary Table 5. List of filtered guide pairs used in the analysis
Supplementary Table 6. Table of observed guide counts in each replicate of the screen
Supplementary Table 7. The locations of functional sequences predicted by RELICS and complete set of RELICS scores for the screened region

#### Supplementary Figures

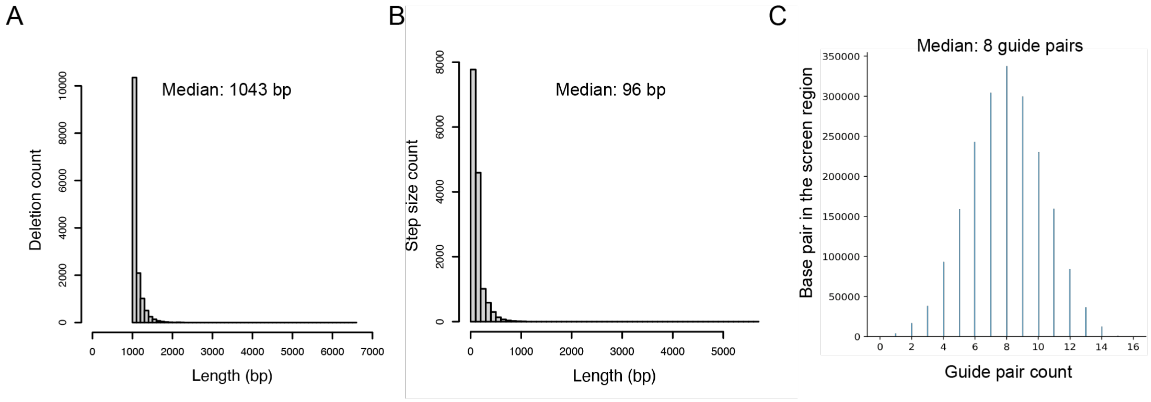


**Supplementary Fig. 1 | The properties of the programmed deletions by 14,555 guide pairs.** (**A**) The distribution of the length of the intended programmed deletions. (**B**) The distribution of the length of the step size between intended programmed deletions. (**C**) Number of intended deletions (guide pairs) that cover each base pair in the screened region.

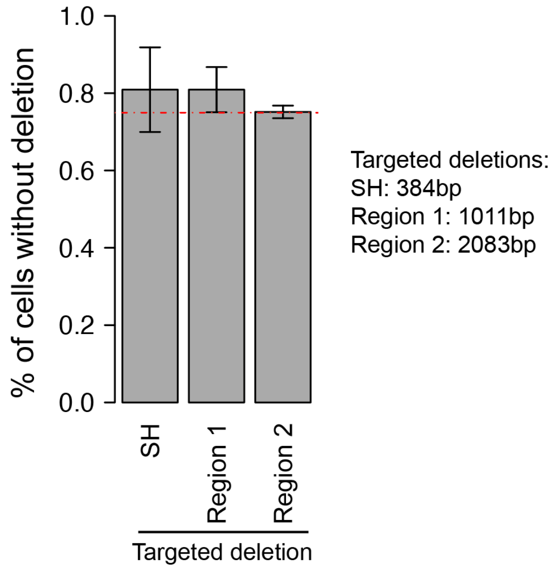


**Supplementary Fig. 2 | Deletion efficiency quantification by qPCR.** qPCR results quantifying the deletion efficiency of three different targeted deletions (Safe Harbor (SH) region, Region 1, or Region 2) in the *GATA3* locus in naïve $CD4^+$ T cells with n=4-6 replicates for each experiment. The red dashed line indicates fold change of 0.75.

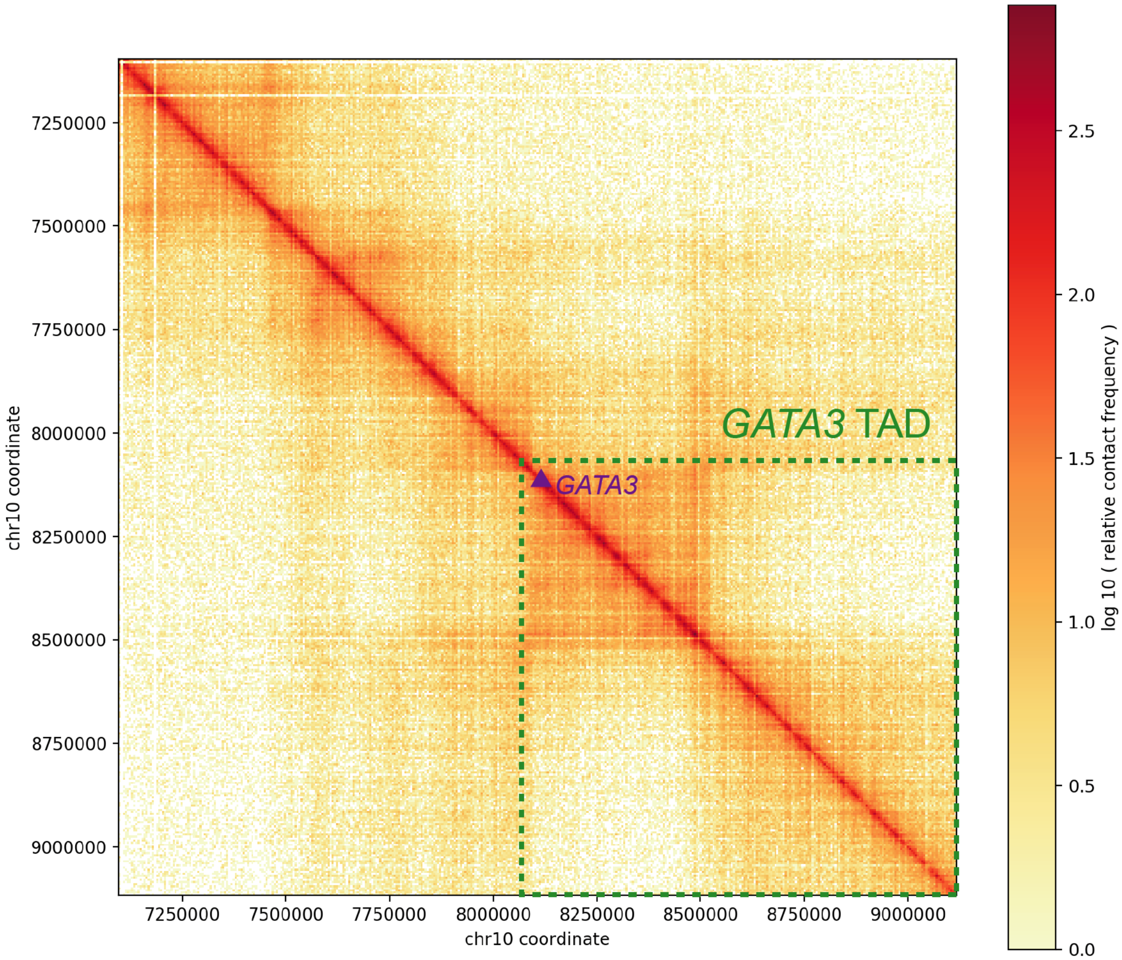


**Supplementary Fig. 3 | Hi-C contact frequency map surrounding *GATA3* in Jurkat cells.** The processed Hi-C data was obtained from Lucic et al. 2019[1] and the region surrounding *GATA3* (chr10:7096666-9117164 (hg19)) was visualized**.** The location of *GATA3* is highlighted in purple triangle and the *GATA3* TAD is indicated with a green square with dashed lines.

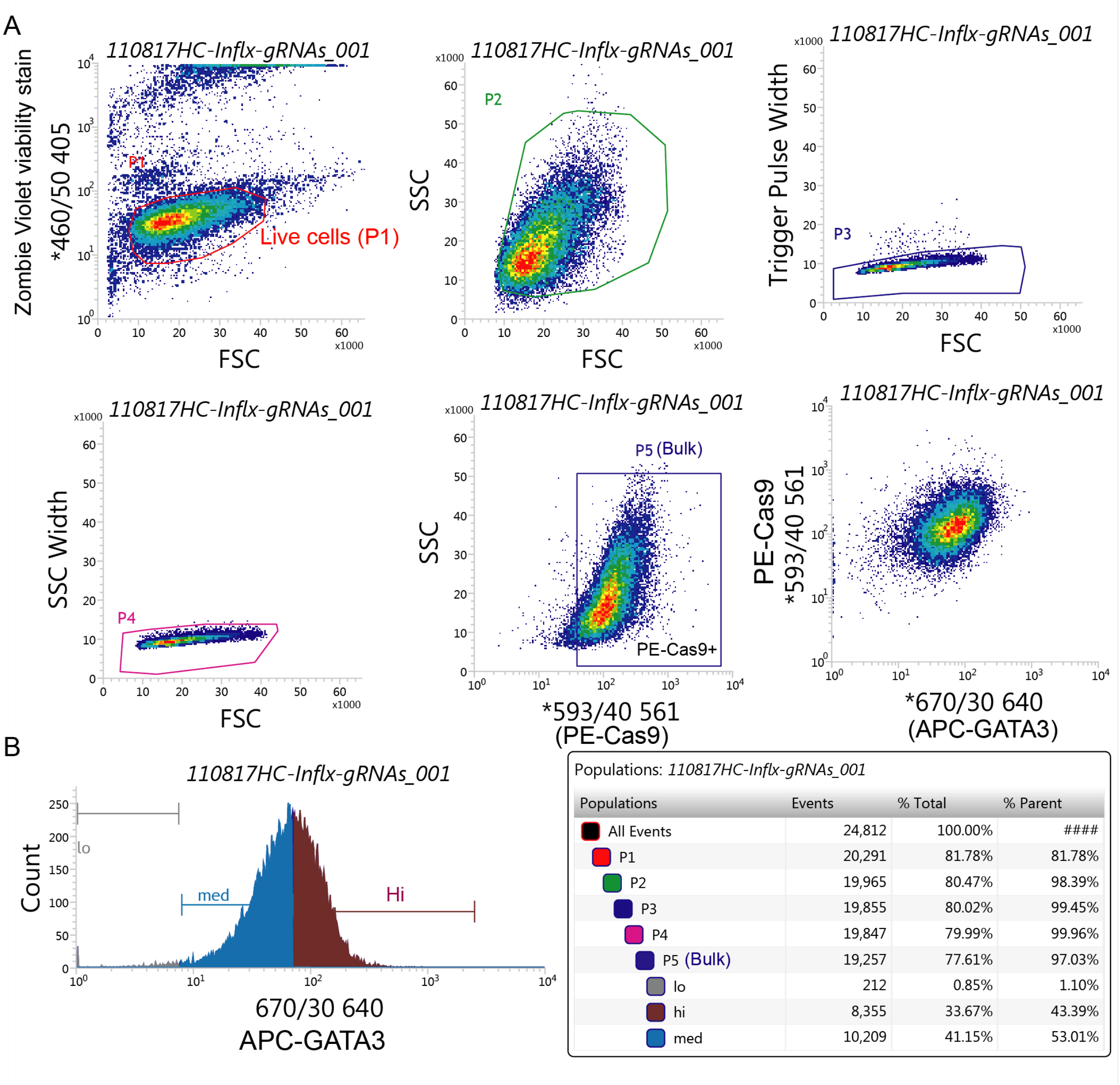


| Populations | Events | % Total | % Parent |
|---|---|---|---|
| All Events | 24,812 | 100.00% | #### |
| P1 | 20,291 | 81.78% | 81.78% |
| P2 | 19,965 | 80.47% | 98.39% |
| P3 | 19,855 | 80.02% | 99.45% |
| P4 | 19,847 | 79.99% | 99.96% |
| P5 (Bulk) | 19,257 | 77.61% | 97.03% |
| lo | 212 | 0.85% | 1.10% |
| hi | 8,355 | 33.67% | 43.39% |
| med | 10,209 | 41.15% | 53.01% |

**Supplementary Fig. 4 | The gating strategy for cell sorting.** (**A**) Data shown is from replicate 1. All events were first gated by the signal from Zombie Violet viability dye staining to obtain live cell population (P1). P1 was then gated by size based on forward scatter (FSC) and side scatter (SSC) to obtain a homogenous population of P4. P4 were further gated by PE-DYKDDDDK-tagged Cas9 to obtain Cas9+ population as bulk (P5). (**B**) The bulk (P5) population was sorted by APC-GATA3 level into low, medium, and high GATA3 expression pools.

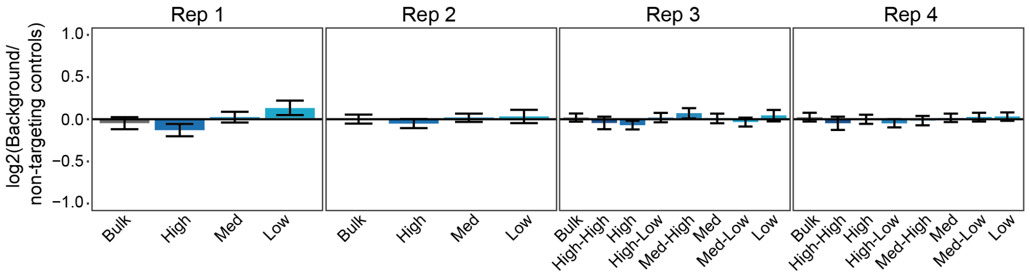


**Supplementary Fig. 5 | Estimated proportion of sgRNA pair counts in each pool in four replicates**. The proportion of sgRNA pair counts targeting outside of *GATA3* gene (background) was compared to non-targeting control sgRNAs in each pool in four replicates. The proportions were estimated by maximum likelihood by RELICS. Whiskers are 95% confidence intervals calculated from 1000 bootstrap iterations.

### GATA3_exon

| Guide Target | PAM Sequence | Indel % | Model Fit ($R^2$) |
|---|---|---|---|
| GGCCCCGGGTGGTGCGTGTCC | GGG | 84 | 0.98 |

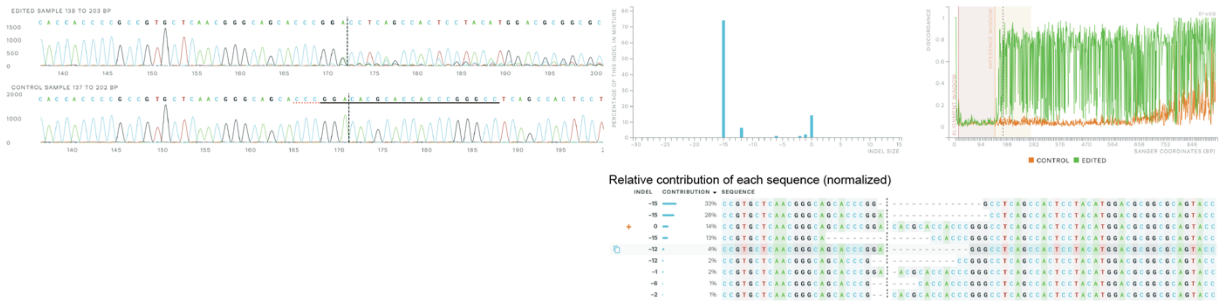


# SH

| Guide Targets | PAM Sequences | Indel % | Model Fit ($R^2$) |
|---|---|---|---|
| g1 GTTGATGCCTTATTACCACC | g1 AGG | 86 | 0.63 |
| g2 CCTGTACTCACACCTACAGC | g2 TGG | | |

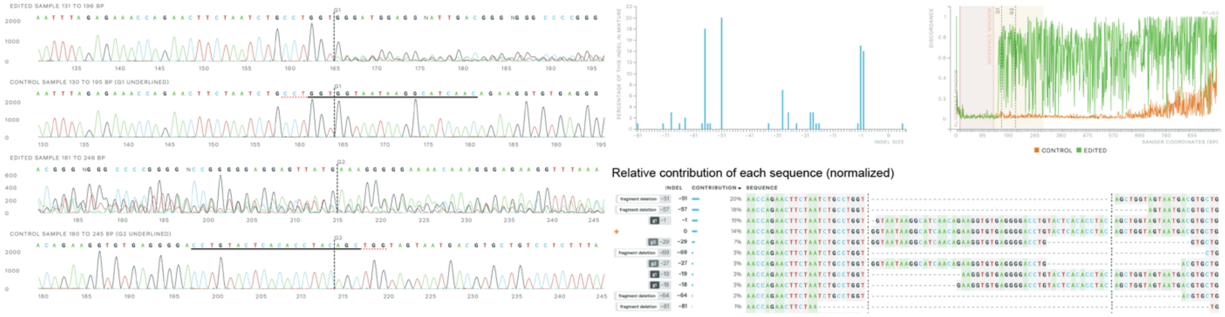


# FS1

| Guide Targets | PAM Sequences | Indel % | Model Fit ($R^2$) |
|---|---|---|---|
| g1 AGAGAGTCAGGGCCTCCATT | g1 AGG | 94 | 0.97 |
| g2 AGGTTGCCATTGCCTCCTAT | g2 GGG | | |

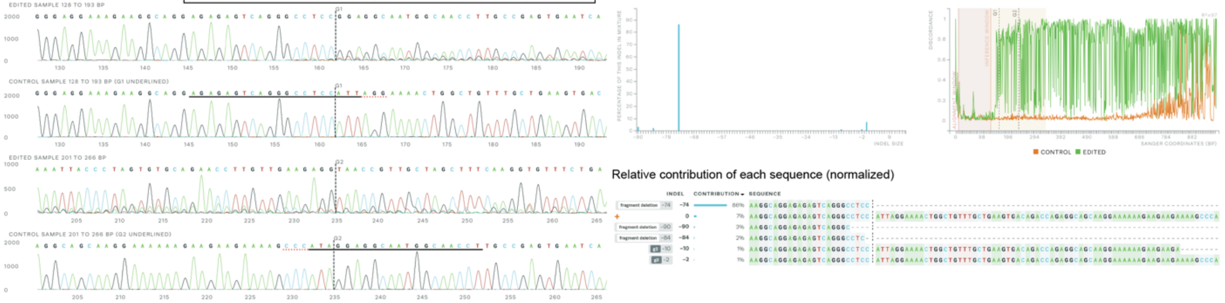


# FS10

| Guide Targets | PAM Sequences | Indel % | Model Fit ($R^2$) |
|---|---|---|---|
| g1 TACTGAGACTGGGTGAGATG | g1 AGG | 98 | 0.95 |
| g2 CACATAGTGAAACCCACATC | g2 AGG | | |

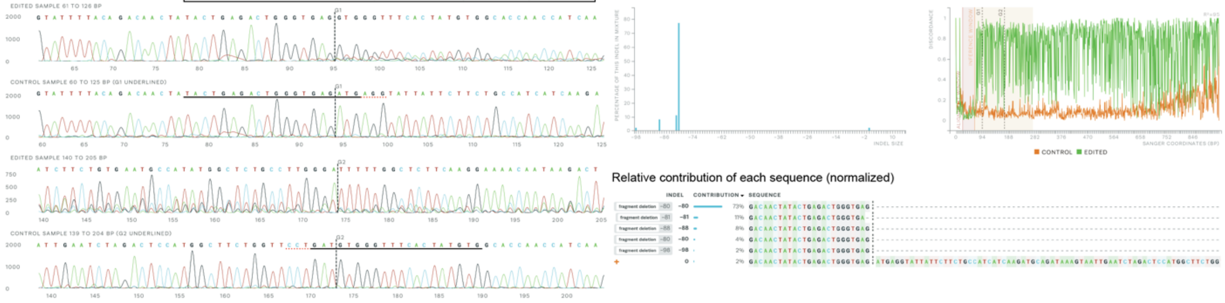

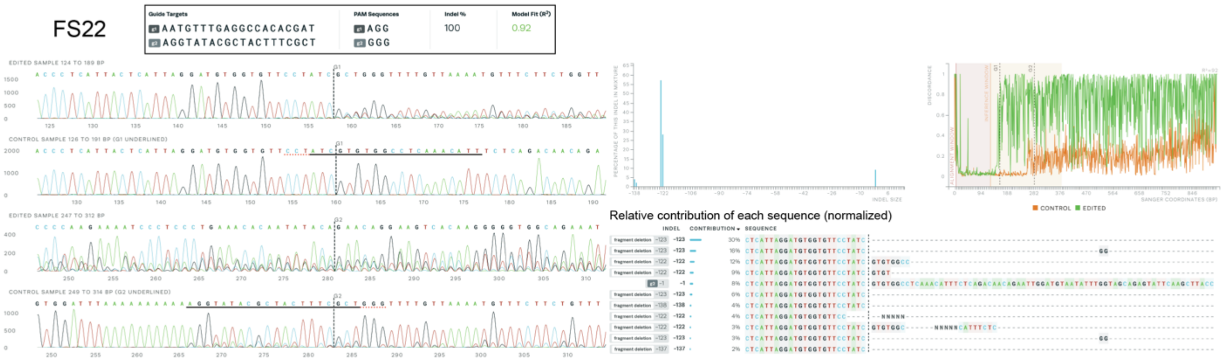


**Supplementary Fig. 6. | Mutation profiling of the CRISPR edited Th2 cells**. Estimated percentage of cells carrying targeted mutation or deletions for *GATA3* exon, safe harbor (SH), FS1, FS10, and FS22 was quantified by tracking of indels by decomposition (TIDE) using the Synthego ICE Analysis tool (Synthego Performance Analysis, ICE Analysis. 2019. v3.0. Synthego). The results are shown in three different plots, the discordance plot which details the level of alignment per base between wild type (control) and the edited sample in the inference window (the region around the cut site), the inferred distribution of indels in the entire edited population of genomes, and the relative contribution of each sequence (normalized). The results for each experiment were reproduced in two independent experiments, one of which is shown.

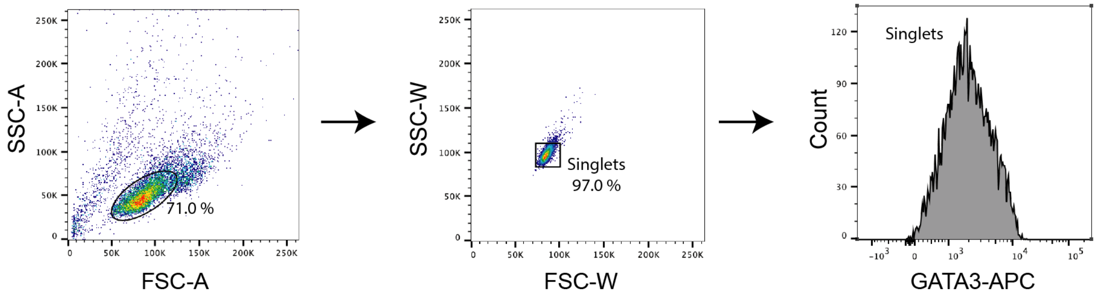


**Supplementary Fig. 7 | A representative diagram showing the gating strategy used for flow cytometry analysis**. All events were first gated by the size of the area of forward scatter (FSC) and side scatter (SSC). The gated population were further gated by the size of the width of FSC and SSC to gain the singlet population for downstream analyses, such as GATA3 expression (APC-A) of the singlets.

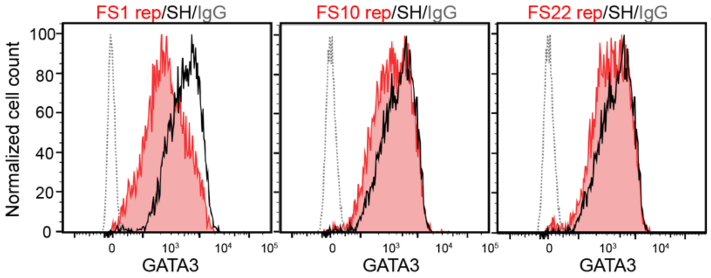


**Supplementary Fig. 8 | Validation of predicted functional sequences in regulating *GATA3* expression in Th2 cells.** Flow cytometry results from intra-cellular staining of GATA3 protein following Cas9 RNP experiments. Each panel also shows results using an isotype control (IgG, dotted line) and from targeting a safe harbor (SH) genome sequence (black outline). These experiments are independent replicates of those experiments shown in Fig. 3c.

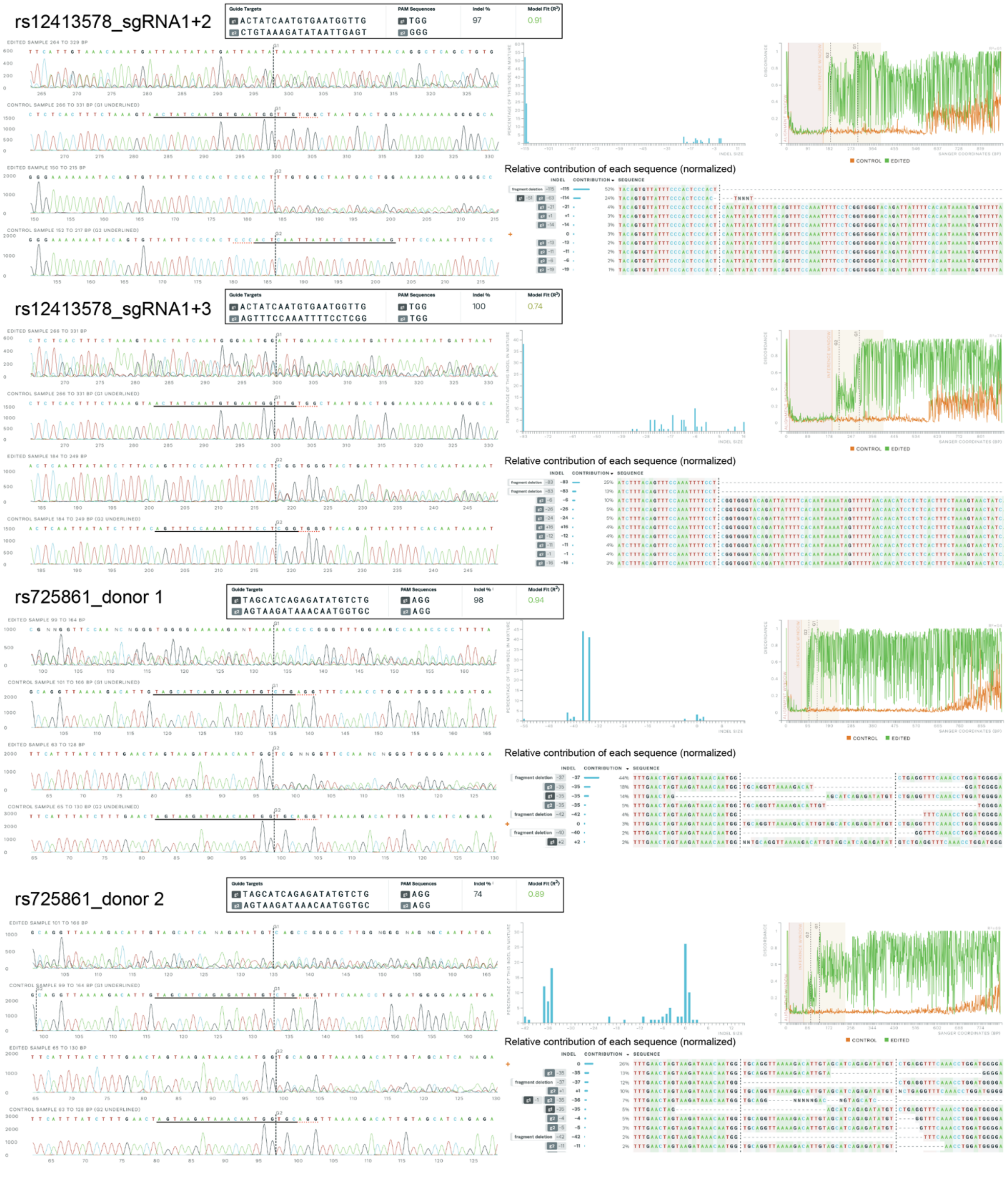


**Supplementary Fig. 9 | Mutation profiling of the CRISPR edited Th2 cells**. Estimated percentage of cells carrying targeted deletions spanning rs12413578 or rs725861 was quantified by tracking of indels by decomposition (TIDE) using the Synthego ICE Analysis tool (Synthego Performance Analysis, ICE Analysis. 2019. v3.0. Synthego). The results are shown in three different plots, the discordance plot which details the level of alignment per base between wild type (control) and the edited sample in the inference window (the region around the cut site), the inferred distribution of indels in the entire edited population of genomes, and the relative contribution of each sequence (normalized). The blood sample used to test rs12413578 was obtained from a donor that carries the risk allele, C, for rs12413578. Both donors used to test rs725861 carry the protective allele, A, for rs725861.

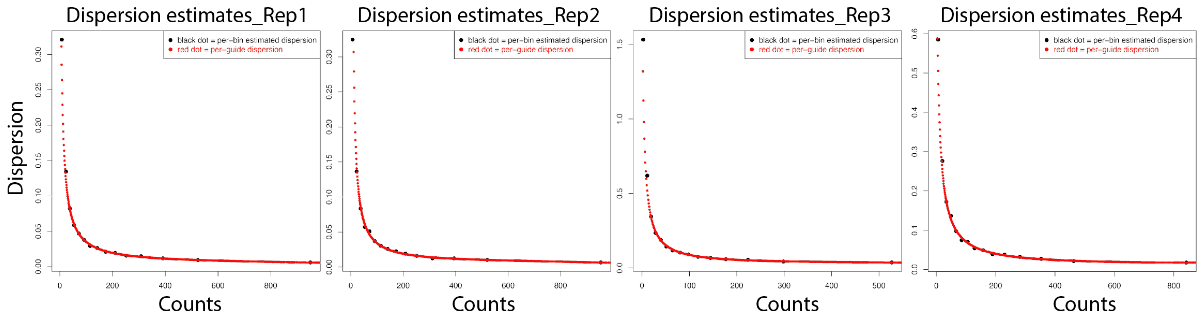


**Supplementary Fig. 10. | Splines with 2 degrees of freedom fit to dispersion estimates binned by total guide counts from RELICS for replicates 1, 2, 3, and 4 of the screen.** The splines are plotted in red and the binned dispersion estimates are indicated with black points.

| Antibody | Manufacture | Catalog number |
|---|---|---|
| APC anti-GATA3 antibody | Biolegend | 653806 |
| Brilliant Violet 510$^{TM}$ anti-human CD4 antibody | Biolegend | 317443 |
| PE anti-DYKDDDDK | Biolegend | 637309 |
| Mouse anti-human IFN-g antibody | BD Pharmingen | 554547 |

**Supplementary Table 2. Antibodies used in this study.**
